# Interactome analysis identifies MSMEI_3879 as a substrate of *Mycolicibacterium smegmatis* ClpC1

**DOI:** 10.1101/2022.10.26.513873

**Authors:** Emmanuel Ogbonna, Priyanka Bheemreddy, Karl R. Schmitz

**Affiliations:** Department of Biological Sciences University of Delaware Newark DE, 19716; Department of Chemistry & Biochemistry University of Delaware Newark DE, 19716

## Abstract

The prevalence of drug resistant *Mycobacterium tuberculosis* infections has prompted extensive efforts to exploit new mycobacterial drug targets. ClpC1, the unfoldase component of the essential ClpC1P1P2 protease, has emerged as one particularly promising antibacterial target. However, efforts to identify and characterize ClpC1-targeting compounds are constrained by our limited knowledge of Clp protease function and regulation. To expand our understanding of ClpC1 physiology, we employed a co-immunoprecipitation and mass spectrometry workflow to identify proteins that interact with ClpC1 in *Mycolicibacterium smegmatis*, a relative of *M. tuberculosis.* We identify a diverse panel of interaction partners, many of which make co-immunoprecipitate with both the regulatory N-terminal domain and the ATPase core of ClpC1. Notably, our interactome analysis identifies MSMEI_3879, a truncated gene product unique to *M. smegmatis*, as a novel proteolytic substrate. Degradation of MSMEI_3879 by ClpC1P1P2 *in vitro* requires an exposed N-terminal sequence, reinforcing the idea that ClpC1 selectively recognizes disordered motifs. Fluorescent substrates incorporating MSMEI_3879 may be useful in screening for novel ClpC1-targeting antibiotics, to help address the challenge of *M. tuberculosis* drug resistance.

## INTRODUCTION

Tuberculosis is responsible for greater global mortality than any other bacterial pathogen, causing an estimated 1.5 million deaths in 2020 alone (1). While rates of infection and mortality have declined over the past decade, the prevalence of multidrug-resistant tuberculosis infections remains a persistent challenge to global public health. There is consequently an urgent need to develop new drugs and exploit new drug targets in the causative pathogen, *Mycobacterium tuberculosis* (*Mtb*). The essential mycobacterial Clp proteases have emerged as one promising class of targets (2–7).

Clp proteases are well studied proteolytic machines that unravel and hydrolyze native protein substrates(8, 9). These large oligomeric complexes consist of a hexameric Clp unfoldase that recognizes substrates, unfolds them using energy from ATP hydrolysis, and spools them into an associated peptidase barrel for degradation (10). The mycobacterial Clp peptidase is composed of two paralogous heptamers, ClpP1 and ClpP2, that assemble into a catalytically active hetero-oligomeric ClpP1P2 tetradecamer (11–16). Two alternative unfoldases, ClpC1 and ClpX, dock on the ClpP2 face of the peptidase to form the functional Clp protease (13, 14, 17, 18). Substrates are recognized by directly interacting with the unfoldase (19–21), with the aid of proteolytic adaptors (22, 23), or via post-translational phosphorylation (24–26).

All components of *M. tubercuosis* Clp proteases are strictly essential for viability (19, 27–31), making these enzymes attractive targets for novel drug development. Multiple classes of antimicrobials inhibit or dysregulate Clp protease activity by targeting the peptidase (2, 5, 13, 14, 29, 32–34). However, several compound classes have been shown to cross-target human mitochondrial CLPP (35–37), which complicates efforts to develop ClpP-targeting pharmacophores into viable antibacterial therapeutics. By contrast, ClpC1 has no direct homologs in animals, and thus provides an avenue for inhibiting Clp protease activity in the pathogen with lower risk of off-target effects in humans. Cyclic peptides have been identified that kill *Mtb* by targeting ClpC1, including cyclomarin A, metamarin, lassomycin, ecumicin and rufomycin, all of which bind to the N-terminal domain (NTD) (3, 4, 38–43).

Bacterial Clp proteases participate in various physiological processes, including protein quality control, response to stress, and regulation of virulence (19, 44–53). Although Clp proteases are known to be essential in mycobacteria, the breadth of their physiological roles are poorly understood. One path to elucidating the functions of Clp proteases is to identify their proteolytic substrates and interaction partners. Prior studies have used targeted capture approaches to identify Clp protease substrates in several bacteria, including *Escherichia coli*, *Bacillus subtilis, Staphylococcus aureus,* and *Caulobacter crescentus* (48, 52, 54–56).

In this study, we use co-immunoprecipitation and mass spectrometry to identify cellular proteins that interact with the ClpC1 unfoldase in *Mycolicibacterium smegmatis* (*Msm*), a nonpathogenic relative of *Mtb*. Importantly, we identify and characterize a novel substrate of the ClpC1P1P2 protease in *Mycolicibacterium smegmatis*.

## Materials and Methods

### Plasmid and strain construction

— Full-length ClpC1 (ClpC1^WT^), the ClpC1 NTD alone (aa 1-147; ClpC1^NTD^), and ClpC1 lacking the NTD (aa 158-848; ClpC1^CORE^) were amplified from *Mycolicibacterium smegmatis* (strain ATCC 700084 / MC^2^155) genomic DNA (ATCC) and cloned into a modified episomal pNIT expression vector (57), followed by a C-terminal 3×FLAG tag. Mutations to Walker B motifs in the D1 (E288Q) and D2 (E626Q) rings of ClpC1^WT^ were introduced by sequential polymerase chain reactions, yielding an ATPase inactive “trap” variant (ClpC1^EQ^) (56). Plasmid sequences were verified by Sanger sequencing (Genewiz).

### In vivo substrate trapping

Plasmids encoding *M. smegmatis* ClpC1 constructs were electroporated into *M. smegmatis* (ATCC 700084 / MC^2^155) at 5 kV in a MicroPulser (Bio-Rad). Liquid starter cultures were made in Middlebrook broth base (HiMedia) supplemented with 0.2% v/v glycerol (Fisher Scientific), 0.2% w/v glucose (TCI), and 0.05% v/v Tween 80, and then (with orbital shaking) grown for 60 hours at 37 °C. At a starting A_600_ of 0.05, starter cultures were sub-cultured into 200 mL of fresh media. The cultures were then grown at 37 °C until mid-log phase (A_600_ ≈ 0.6-1.0), at which point expression was induced by 28 mM ε-caprolactam (Sigma-Aldrich). For cell harvesting, centrifugation was carried out at 9,000×g for 20 minutes at 4 °C and pellets resuspended in 5 mL of Lysis Buffer (25 mM HEPES, 10 mM magnesium chloride, 200 mM potassium chloride, 0.1 mM EDTA; supplemented with 10 mM ATP, pH 7.5). Cell lysis was done in a microfluidizer (Microfluidics), and subsequent clarification of lysates was at 16,000×g for 30 minutes at 4 °C. Cell supernatants were stored at −80 °C. Bradford assay (Bio-Rad) was performed to estimate total protein content.

### Co-immunoprecipitation

40 µL of EZView Red ANTI-FLAG M2-Affinity Gel beads (Sigma-Aldrich) were equilibrated and washed twice in 0.5 mL Lysis Buffer by centrifugation at 8,200×g for 30 s. To pull down the expressed 3×FLAG-tagged *M. smegmatis* ClpC1 constructs and interacting proteins, the equilibrated beads were incubated with 1 mL of lysate for 1 hour at 4 °C, with gentle agitation. After incubation, the bead-lysate slurry was spun at 8,200×g for 30 s and the bead pellet subsequently washed with Lysis Buffer (containing 10 mM ATP). To elute, the washed beads were then mixed with 20 µL of 2× Laemmli Sample Buffer (10% glycerol, 4% SDS, 167 mM Tris-HCl, 0.02% bromophenol blue) in the absence of reducing agent and boiled for 5 min. Samples were then vortexed briefly and centrifuged, and the supernatant containing the eluate was stored at −80 °C prior to further analysis. As a negative control, lysates of cells containing empty pNIT vector were processed by the same workflow.

### Mass spectrometry sample preparation

Proteins in the eluate were analyzed on a 6-15% SDS-PAGE gradient gel. Each lane was diced, and gel pieces put into 1.5 mL microcentrifuge tubes. The samples were prepared by adapting standard protocols for in-gel mass spectrometry sample preparation (58). Incubation with 10 mM dithiothreitol (DTT) was carried out for 45 min at 55 °C in order to reduce disulfide bonds. Afterwards, carbamidomethylation of cysteines was with 55 µM iodoacetamide for 30 min at room temperature (in the dark). Gel pieces were washed with Gel Wash Buffer (25 mM ammonium bicarbonate and 50% acetonitrile), and dehydrated with 100% acetonitrile prior to drying in a SpeedVac vacuum centrifuge (Thermo). Trypsin digestion was carried out in 25 mM ammonium bicarbonate containing 10 µg ml^−1^ MS Grade Trypsin Protease (Pierce), and incubated overnight at 37 °C. Peptides were subsequently extracted by a two-step process which was repeated twice: incubation of gel pieces with 5% formic acid (Sigma-Aldrich) followed by 100% acetonitrile. To remove residual salts, the peptide samples were desalted using a HyperSep C18 column (Thermo Fisher Scientific) according to previously described procedure (58, 59). C18 column-eluted samples were dried by vacuum centrifugation and frozen prior to mass spectrometry.

### LC-MS/MS

— Liquid chromatography tandem mass spectrometry (LC-MS/MS) analysis was performed using a Q Exactive Orbitrap interfaced with Ultimate 3000 Nano-LC system (Thermo Fisher Scientific). Trypsin-digested samples were loaded on an Acclaim PepMap RSLC column (75 µm x 15 cm nanoViper) using an autosampler. Analysis of samples was done using a 150-min gradient running from 2 % to 95 % Buffer B (0.1 % formic acid in acetonitrile) in Buffer A (0.1% formic acid in water) at a flow rate of 0.3 µL min^−1^. MS data acquisition was done using a data-dependent top10 method, with the most abundant precursor ions from the survey scan chosen for higher energy collisional dissociation (HCD) fragmentation using a stepped normalized collision energy of 28, 30, and 35 eV. Survey scans were acquired at a resolution of 70,000 at m/z 200 on the Q Exactive. LS-MS/MS data was collected in independent biological triplicates.

### MS Data Analysis

For proteomic analysis, extraction of raw data was performed in the Proteome Discoverer software suite (version 1.4, Thermo Fisher Scientific). The raw data was searched against the *Mycolicibacterium smegmatis* (strain ATCC 700084 / MC^2^ 155) UniProt Reference Proteome (Proteome ID UP000000757) using Sequest HT (University of Washington and Thermo Fisher Scientific). Iodoacetamide-mediated cysteine carbamidomethylation was set as a static modification. Precursor mass tolerance was set at 10 ppm, while allowing for fragment ion mass deviation of 0.6 Da for the HCD data, and full trypsinization with a maximum of two missed cleavages. Peptide-spectrum match (PSM) validation was done using Percolator with false discovery rates (FDR) of 1% and 5% for stringent and relaxed validation, respectively. Gene ontology (GO) annotation analysis on the datasets was performed using the Blast2GO software suite (60).

### Expression and purification of recombinant proteins

N-terminal H_7_-SUMO-tagged ClpC1 (with or without a C-terminal sortase tag) and C-terminal H_6_-tagged ClpP1 and ClpP2 were cloning into pET22b-derived vectors using Gibson Assembly (61). N-terminal H_7_-SUMO-tagged putative substrates and interaction partners were obtained as synthetic gene constructs (Twist Bioscience) and cloned into pET29b(+). All constructs were expressed in *E. coli* ER2566 (NEB). Cultures were grown in 1.5×YT at 37 °C to exponential phase (A_600_ 0.8-1.0), and overexpression was induced with 0.5 mM IPTG followed by incubation at 30 °C for 4 h. Cells were harvested at 4,000 ×g for 30 min and pellets resuspended in 25 mL His-tag Lysis Buffer (25mM HEPES pH 7.5, 300mM NaCl, 10mM imidazole pH 7.5, 10% glycerol), supplemented with 1 mM PMSF and 100 µL of EDTA-free protease inhibitor cocktail (Thermo Fisher). After sonication and clarification at 15,000×g for 30 min, lysates were loaded onto a Ni-NTA column (MCLAB), washed with 25 mM imidazole and eluted in 300 mM imidazole. Eluate was spin-concentrated (10,000 MWCO Amicon, MilliporeSigma) at 4,000×g. Protein samples were further purified by anion-exchange chromatography (Source 15Q 10/100, Cytiva). H_7_-SUMO tags were removed by incubation with *Saccharomyces cerevisiae* SUMO protease Ulp1 (62) prior to gel filtration (HiLoad 16/600 Superdex 200, Cytiva) into Protein Degradation Buffer (25 mM HEPES, 200 mM potassium chloride, 10 mM magnesium chloride, 0.1 mM EDTA, pH 7.5).

### ClpC1P1P2 in vitro degradation assays

*In vitro* degradation assays containing 1 µM ClpC1 (hexamer), 1 µM ClpP1 (tetradecamer), 1 µM ClpP2 (tetradecamer) and 10 µM substrate were performed in ClpC1 protein degradation buffer. All degradation assays were carried out in the presence of 50 µM activator peptide Z-Leu-Leu-Nva-CHO (benzyloxycarbonyl-L-leucyl-L-leucyl-L-norvalinal) (12, 13), with a total of 15 mM ATP, along with an ATP regeneration system consisting of 187.5 U mL^−1^ pyruvate kinase and 50 mM phosphoenolpyruvate (Sigma). For gel degradation assays, 14 µL aliquots were taken at each time point and mixed with 7 µL 2× Laemmli Sample Buffer (containing 10% β-mercaptoethanol), and analyzed by SDS-PAGE. Gels were stained by 0.1% Coomassie Brilliant Blue and quantified by ImageJ (63). For plate reader assays, degradation of GFP-substrate fusions was monitored by loss of 511 nm emission following excitation at 450 nm.

### Data availability

The mass spectrometry data from this work have been submitted to the ProteomeXchange Consortium via the PRIDE partner repository (64), and assigned the identifier PXD030385.

## Results

### Identification of interaction partners of full-length ClpC1

We sought to design a set of ClpC1 constructs that maximized our likelihood of identifying diverse interactions partners. Some interaction partners likely bind to wild-type ClpC1 (ClpC1^WT^) irrespective of its nucleotide-bound state; for example, to the flexible NTD. However, substrate binding to wild-type AAA+ unfoldases is often transient, due to unfolding/degradation, and ATP-dependent (8, 9, 65–67). To increase our ability to capture ATP-dependent interactions and prevent degradation of *bona fide* substrates, we mutated conserved Glu residues within the D1 and D2 Walker B motifs to Gln (E288Q and E626Q, respectively), yielding ClpC1^EQ^ (**Figs. 1A, B**) (68). Analogous mutations in related AAA+ enzymes stabilize ATP binding but impair nucleotide hydrolysis and substrate unfolding (56, 69, 70). To facilitate co-immunoprecipitation with interaction partners, all constructs incorporated a C-terminal 3×FLAG tag (DYKDHDG-DYKDHDI-DYKDDDDK).

**Figure 1:**
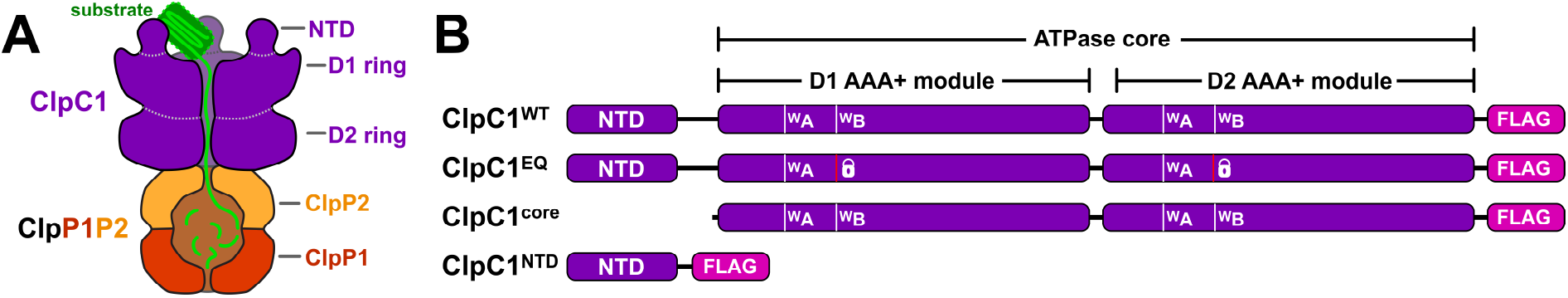
ClpC1 Walker B double mutant substrate trap was generated in *Mycolicibacterium smegmatis* (strain ATCC 700084 / MC^2^155). **(A)** General architecture of the mycobacterial Clp protease. The ATP-dependent unfoldase ClpC1 is responsible for recognizing (via ^ClpC1^NTD) and unfolding protein substrates in an ATP-dependent manner (via AAA+ domains ^ClpC1^D1 and ^ClpC1^D2), and translocating them into the peptidase barrel composed of catalytic heptamers ClpP1 and ClpP2, wherein degradation occurs. ClpC1^EQ^ associates stably with protein substrates but is unable to unfold or translocate them. **(B)** Linear sequence models showing the ClpC1 constructs used in this study to identify cellular proteins interacting with specific ClpC1 components such as the NTD and AAA+ core.

ClpC1 constructs, or an empty vector control, were expressed in *M. smegmatis* MC^2^ 155. ClpC1 and interaction partners were co-immunoprecipitated from lysates in the presence of 10 mM ATP using α-FLAG beads and separated by SDS-PAGE (**Fig. 2**). Samples were prepared by in-gel trypsin digestion and analyzed by LC-MS/MS using higher energy collisional dissociation (HCD) fragmentation. Proteomic analysis was performed in the Proteome Discoverer software suite against the *Mycolicibacterium smegmatis* MC^2^155 proteome using Sequest HT. Proteins observed in at least two of three biological replicates were considered for further analysis. Five proteins were observed in the empty vector control (**Fig. 2**), presumably due to nonspecific interactions with beads. Occurrences of these proteins were ignored in all other samples. In total, 163 and 93 proteins were identified in ClpC1^WT^ and ClpC1^EQ^ datasets, respectively (**Fig. 3A; Supplemental Tables S1, S2**).

**Figure 2:**
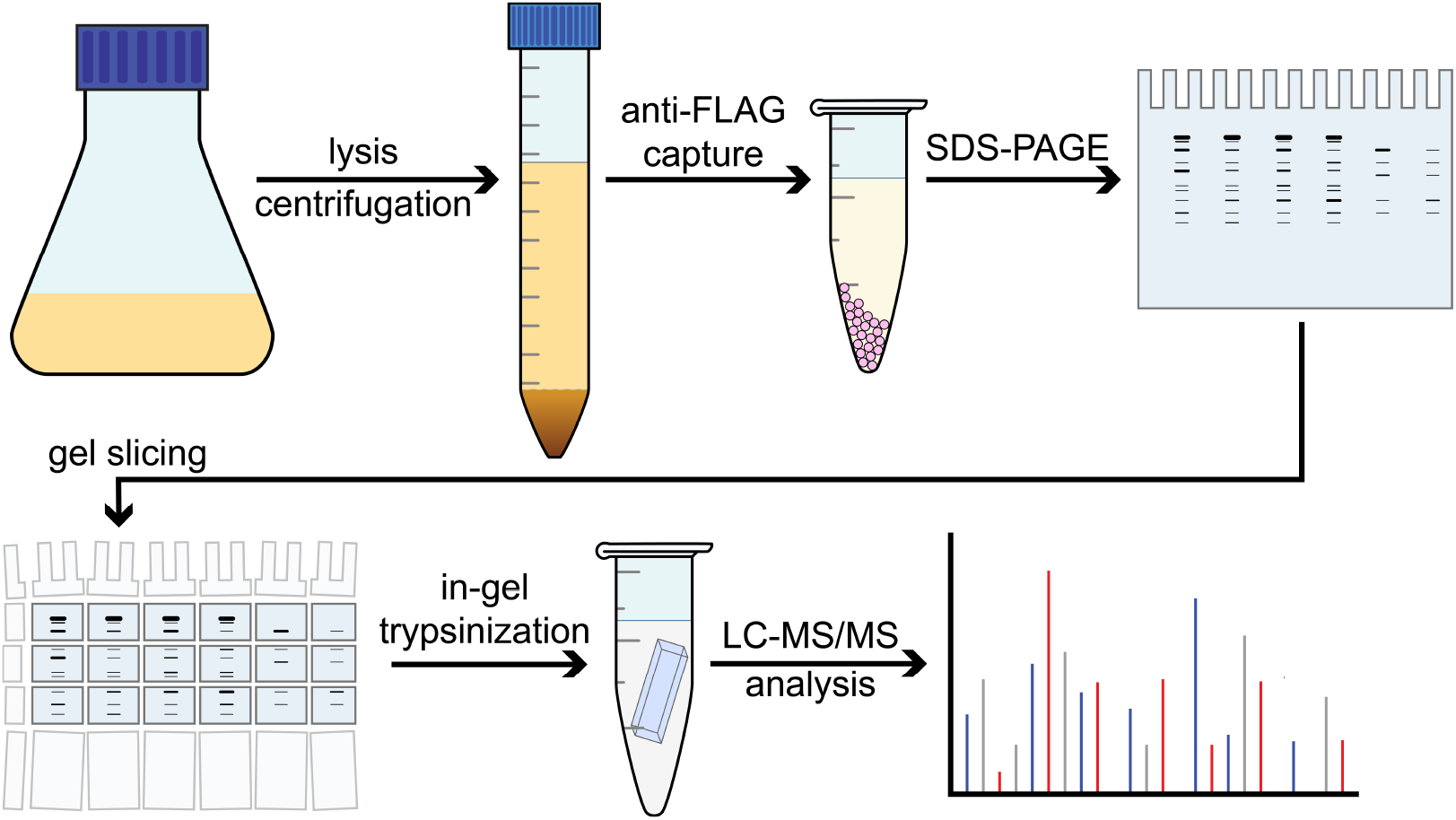
Workflow for identification of ClpC1 interaction partners. *Mycolicibacterium smegmatis strain* (ATCC 700084 / MC^2^155) cells are grown up to A_600_ ≈ 0.6-1.0, and ClpC1 expression is induced following addition of 28 mM ε-caprolactam. Identification of the interactome was based on gel electrophoresis, trypsin digestion, and LC-MS/MS run using higher energy collisional dissociation (HCD) fragmentation. All ClpC1 test construct and control experiments were subjected to the same workflow, and were carried out in biological replicates.

**Figure 3:**
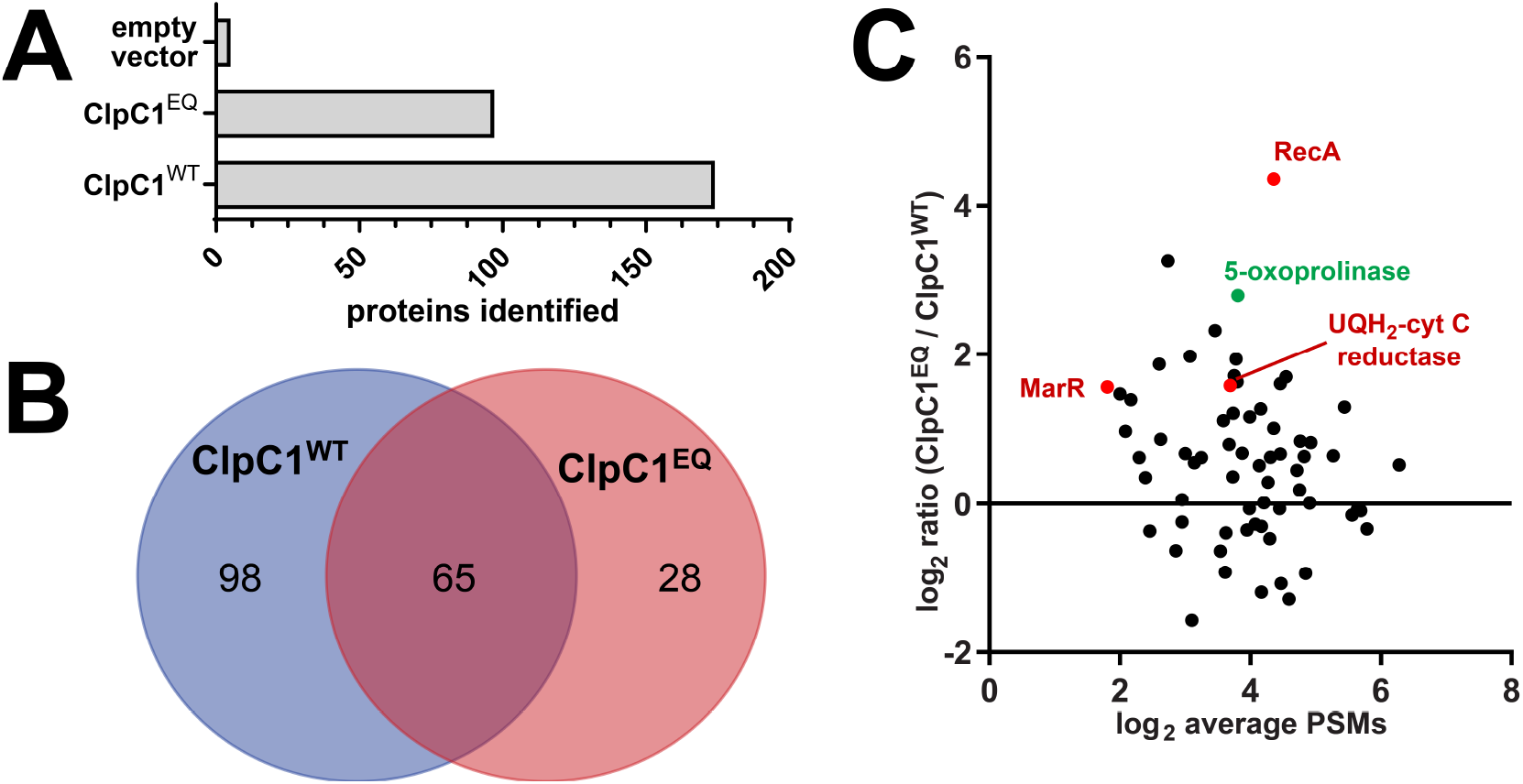
Comparative analysis between ClpC1^EQ^ and ClpC1^WT^ interactome. **(A)** Bar graph showing the number of proteins identified in the ClpC1^EQ^ and ClpC1^WT^ datasets. **(B)** Venn diagram showing the number of *Mycolicibacterium smegmatis* cellular proteins that interacted with ClpC1^EQ^ and ClpC1^wt^constructs, respectively. The numbers in Figs. **A** and **B** exclude ClpC1 protein itself. **(C)** Volcano plot showing enrichment of proteins captured in *Msm* ClpC1 mutant trap versus wild type by liquid chromatography-tandem mass spectrometry (LC-MS/MS). *Y*-axis shows the log2 ratio of normalized scores in the ClpC1^EQ^ dataset to those in the ClpC1^WT^ dataset, while *X*-axis shows log2 average number of PSMs for the proteins. Red circles represent proteins enriched in ClpC1^EQ^ with a p value<0.05 upon Student’s t-test.

We expected substrates to bind more stably to ClpC1^EQ^ than to ClpC1^WT^ because the wild-type enzyme can presumably unfold substrates in the pulldown conditions. Within the ClpC1^EQ^ dataset, 23 proteins matched these criteria and were absent from the ClpC1^WT^ data (**Fig. 3A, Table 1**). These include cell wall synthesis protein Wag31, succinate dehydrogenase iron-sulfur protein SdhB, aconitate hydratase A (AcnA), and ATP synthase epsilon chain AtpC. There were 65 proteins observed in both the ClpC1^WT^ and ClpC1^EQ^ datasets (**Fig. 3A and B**). Of these, 20 were enriched in ClpC1^EQ^ over ClpC1^WT^ by at least a ratio of 2:1 in average normalized score (**Fig. 3B**). However, only three proteins were enriched significantly, with a p-value < 0.05: the DNA repair and stress response protein RecA; electron transport chain component Ubiquinol-cytochrome c reductase cytochrome b subunit (MSMEG_4263); and a MarR-family protein transcriptional regulator (MSMEG_2538) (**Fig 3B**). The co-occurrence of these in ClpC1^WT^ and ClpC1^EQ^ datasets suggests that these interactions occur stably regardless of ATPase and unfolding activity, as would be expected for adaptors, regulators, or other non-substrate interactors.

**Table 1.** Proteins that co-immunoprecipitate with ClpC1EQ but not ClpC1WT.

| Uniprot Accession | Gene Name | Protein | Normalized Average Score* | Average #PSMs |
| --- | --- | --- | --- | --- |
| A0R461 | MSMEG_5715 | Bac_luciferase domain-containing protein | 0.390 | 10.667 |
| A0QWT6 | MSMEG_3058 | Lipoprotein | 0.144 | 3.000 |
| A0QSP8 | rplM | 50S ribosomal protein L13 | 0.085 | 6.667 |
| A0QSD2 | rplD | 50S ribosomal protein L4 | 0.048 | 5.000 |
| A0QRD2 | MSMEG_1073 | Oxidoreductase, short-chain dehydrogenase/reductase family protein | 0.091 | 9.333 |
| A0R1Z9 | atpC | ATP synthase epsilon chain (ATP synthase F1 sector epsilon subunit) | 0.060 | 3.000 |
| A0QVU2 | MSMEG_2695 | 35 kDa protein | 0.083 | 2.500 |
| A0QWK5 | MSMEG_2974 | Peptidyl-prolyl cis-trans isomerase (PPIase) | 0.085 | 2.500 |
| A0QZ33 | MSMEG_3880 | Nitrilase/cyanide hydratase and apolipoprotein N-acyltransferase | 0.130 | 17.500 |
| A0R1H2 | MSMEG_4752 | Uncharacterized protein | 0.062 | 3.333 |
| A0R5K8 | MSMEG_6227 | Transcriptional regulator, PadR family protein | 0.057 | 8.000 |
| A0R006 | wag31 | Cell wall synthesis protein Wag31 (Antigen 84) | 0.075 | 5.667 |
| A0R2B0 | MSMEG_5048 | Uncharacterized protein | 0.035 | 5.500 |
| A0QPX3 | MSMEG_0550 | Sulfonate binding protein | 0.085 | 3.500 |
| A0QSG6 | rpsE | 30S ribosomal protein S5 | 0.075 | 12.333 |
| A0R1B5 | MSMEG_4692 | Uncharacterized protein<br>MSMEG_4692/MSMEI_4575 | 0.082 | 4.500 |
| A0QTM5 | pcaD | 3-oxoadipate enol-lactonase (EC 3.1.1.24) | 0.102 | 4.500 |
| A0QU11 | MSMEG_2037 | Bac_luciferase domain-containing protein | 0.030 | 3.000 |
| A0QS62 | rplJ | 50S ribosomal protein L10 | 0.078 | 8.500 |
| A0QX20 | acnA | Aconitate hydratase A (ACN) (Aconitase) (EC 4.2.1.3) | 0.055 | 14.500 |
| A0QR90 | MSMEG_1029 | Probable transcriptional regulatory protein | 0.095 | 7.500 |
| A0QT07 | sdhB | Succinate dehydrogenase, iron-sulfur protein (EC 1.3.99.1) | 0.059 | 6.500 |
| A0QS46 | rplA | 50S ribosomal protein L1 | 0.053 | 12.000 |
| A0QZJ0 | MSMEG_4042 | Transcriptional regulator, GntR family protein | 0.050 | 6.000 |
| A0QS97 | rpsG | 30S ribosomal protein S7 | 0.082 | 14.500 |
| Q3L885 | pks | Polyketide synthase (Type I modular polyketide synthase) | 0.023 | 43.000 |
| A0QSG5 | rplR | 50S ribosomal protein L18 | 0.032 | 6.500 |
| A0R616 | MSMEG_6391 | Propionyl-CoA carboxylase beta chain (EC 6.4.1.3) | 0.039 | 9.500 |

\* Normalized score is the percentage of the total ion score of a single run attributable to a given protein. Normalized scores were averaged across replicates.

We additionally cross-referenced the 191 proteins observed across ClpC1^WT^ and ClpC1^EQ^ datasets with *Mtb* Clp protease interaction partners identified elsewhere by bacterial two-hybrid screening (23) or by LC-MS/MS upon knockdown of ClpP1 and ClpP2 (21). Thirty *M. smegmatis* proteins in our full-length ClpC1 interactome have *M. tuberculosis* orthologs identified in these prior studies (**Table 3**).

**Table 2.**
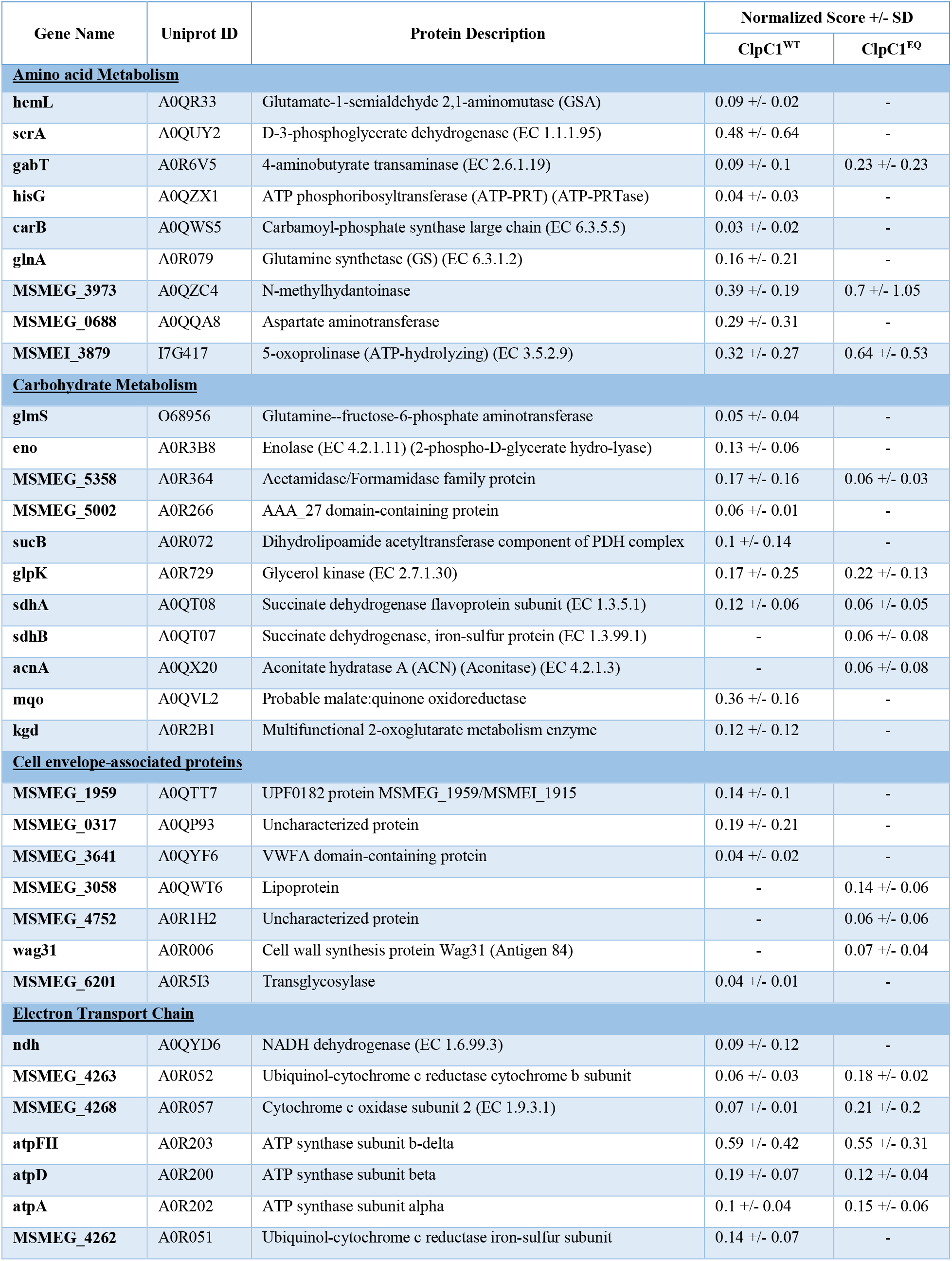
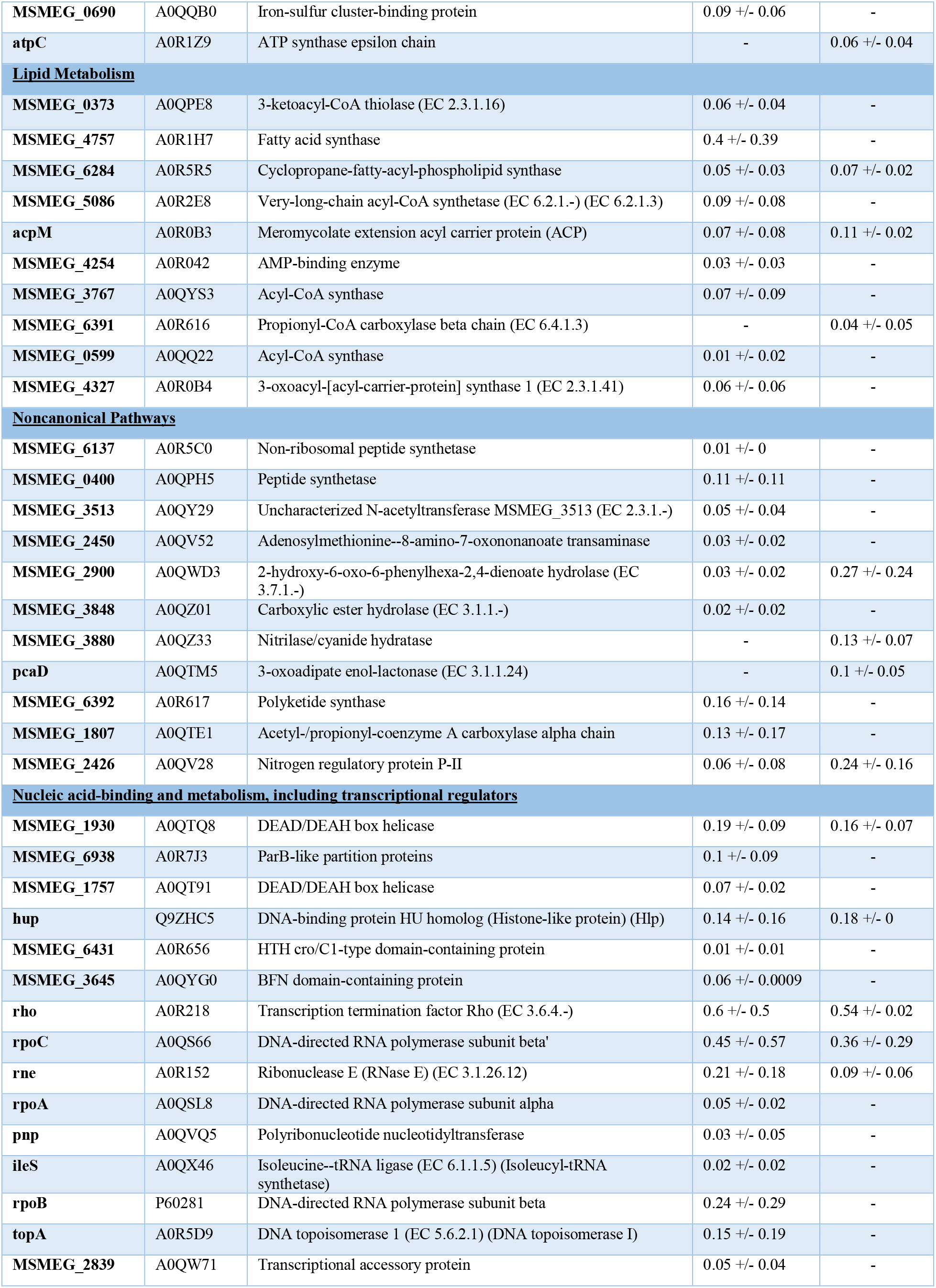
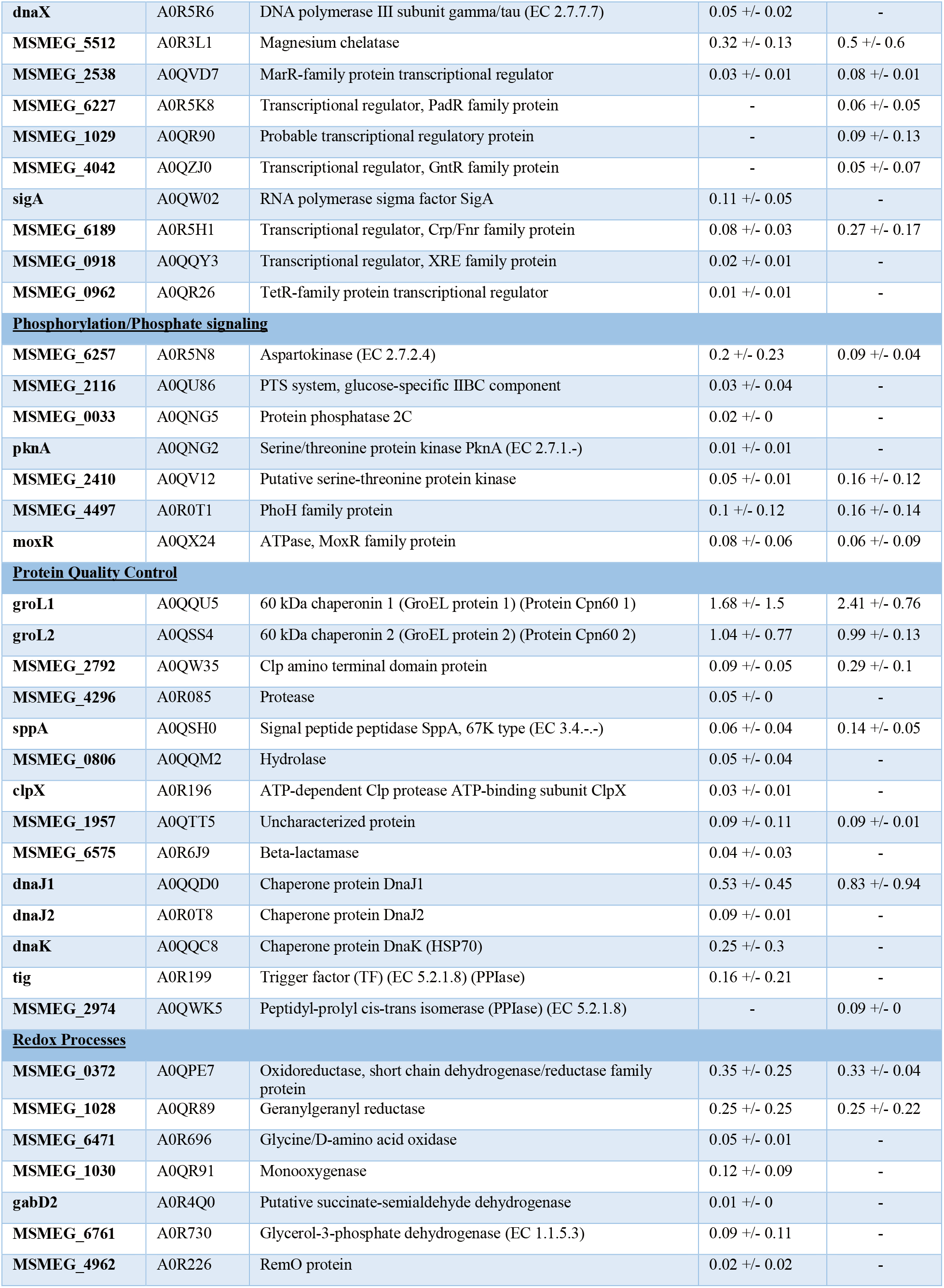
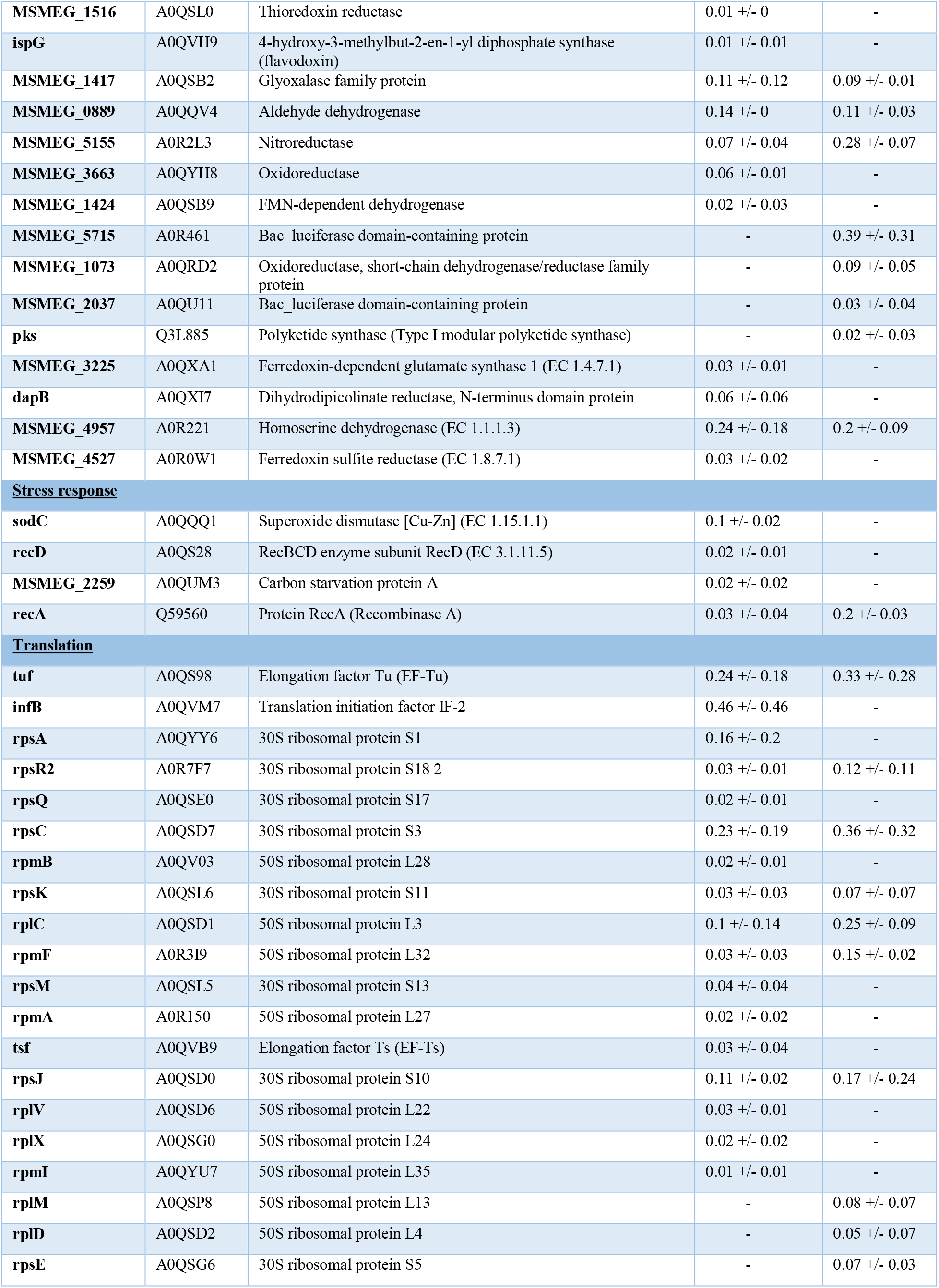
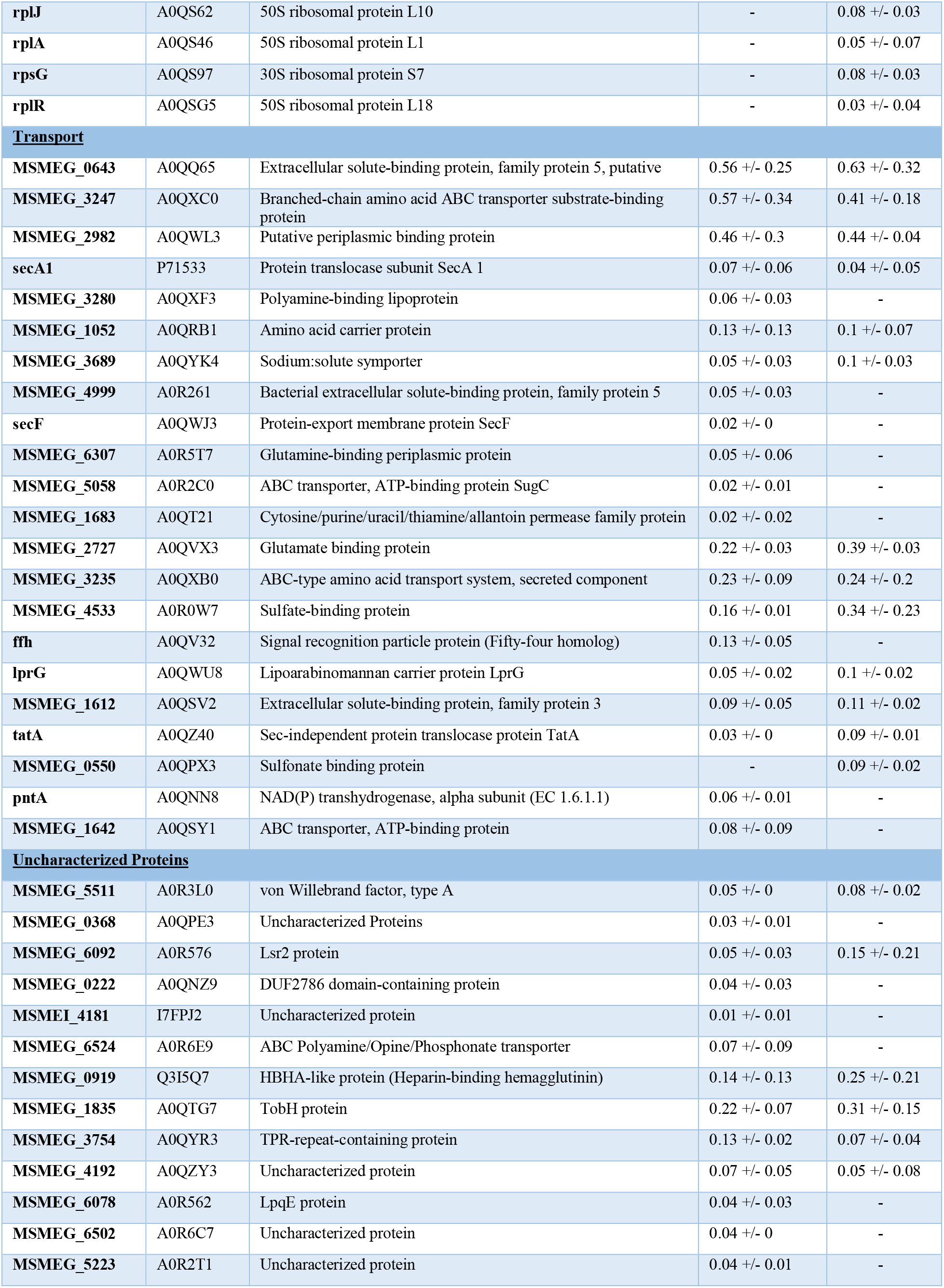
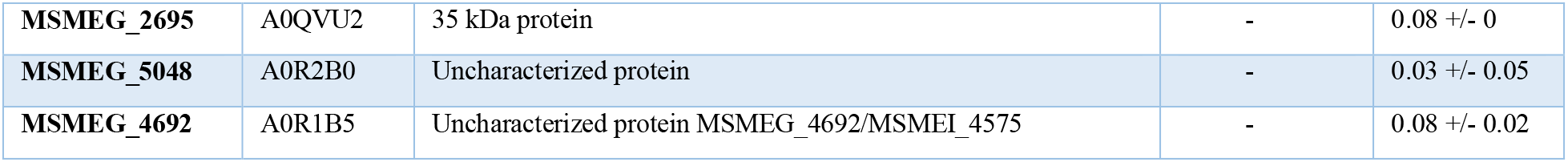
Proteins that interact with full-length *Msm*ClpC1 (ClpC1wt or ClpC1EQ).

**Table 3:** ClpC1 interaction partners for which *M. tuberculosis* orthologs were identified in prior ClpC1 interaction studies.

| Uniprot Accession | Gene name | Protein Description | Prior Work(s) | Mean Norm. Score | Mean #PSMs |
| --- | --- | --- | --- | --- | --- |
| A0QQD0 | dnaJ1 | Chaperone protein DnaJ | ClpC1/P2 KD <sup>1</sup> | 0.676 | 38.500 |
| A0R203 | atpFH | ATP synthase subunit b-delta | ClpC1/P2 KD <sup>1</sup> | 0.568 | 51.667 |
| I7G417 | MSMEI_3879 | 5-oxoprolinase | ClpC1/P2 KD <sup>1</sup> | 0.480 | 20.500 |
| A0R3L1 | MSMEG_5512 | Magnesium chelatase | ClpC1/P2 KD <sup>1</sup>, BACTH <sup>2</sup> | 0.410 | 28.333 |
| A0QQA8 | MSMEG_0688 | Aspartate aminotransferase | ClpC1/P2 KD <sup>1</sup> | 0.292 | 30.500 |
| A0R079 | glnA | Glutamine synthetase (GS) | ClpC1/P2 KD <sup>1</sup> | 0.162 | 14.000 |
| A0R617 | MSMEG_6392 | Polyketide synthase | ClpC1/P2 KD <sup>1</sup> | 0.160 | 35.333 |
| A0R200 | atpD | ATP synthase subunit beta | ClpC1/P2 KD <sup>1</sup> | 0.158 | 11.667 |
| A0R0T1 | MSMEG_4497 | PhoH family protein | BACTH <sup>2</sup> | 0.129 | 9.500 |
| A0QV32 | ffh | Signal recognition particle protein (Fifty-four homolog) | ClpC1/P2 KD <sup>1</sup> | 0.126 | 39.500 |
| A0R202 | atpA | ATP synthase subunit alpha | ClpC1/P2 KD <sup>1</sup> | 0.124 | 14.667 |
| A0QQV4 | MSMEG_0889 | Aldehyde dehydrogenase | ClpC1/P2 KD <sup>1</sup> | 0.123 | 15.417 |
| A0QQQ1 | sodC | Superoxide dismutase | ClpC1/P2 KD <sup>1</sup> | 0.100 | 6.500 |
| A0R0T8 | dnaJ2 | Chaperone protein DnaJ | ClpC1/P2 KD <sup>1</sup> | 0.094 | 21.000 |
| A0QYD6 | ndh | NADH dehydrogenase (EC 1.6.99.3) | ClpC1/P2 KD <sup>1</sup> | 0.093 | 27.333 |
| A0QR33 | hemL | Glutamate-1-semialdehyde 2,1-aminomutase (GSA) | ClpC1/P2 KD <sup>1</sup> | 0.086 | 18.500 |
| A0QVU2 | MSMEG_2695 | 35 kDa protein | ClpC1/P2 KD <sup>1</sup> | 0.083 | 2.500 |
| A0QSY1 | MSMEG_1642 | ABC transporter, ATP-binding protein | ClpC1/P2 KD <sup>1</sup>, BACTH <sup>2</sup> | 0.083 | 30.500 |
| A0QX24 | moxR | ATPase, MoxR family protein | ClpC1/P2 KD <sup>1</sup> | 0.073 | 5.500 |
| A0QZ40 | tatA | Sec-independent protein translocase protein TatA | ClpC1/P2 KD <sup>1</sup> | 0.060 | 4.500 |
| A0R1Z9 | atpC | ATP synthase epsilon chain | ClpC1/P2 KD <sup>1</sup> | 0.060 | 3.000 |
| A0QT07 | sdhB | Succinate dehydrogenase, iron-sulfur protein | ClpC1/P2 KD <sup>1</sup> | 0.059 | 6.500 |
| A0QNN8 | pntA | NAD(P) transhydrogenase, alpha subunit (EC 1.6.1.1) | BACTH <sup>2</sup> | 0.059 | 18.000 |
| P71533 | secA1 | Protein translocase subunit SecA 1 | ClpC1/P2 KD <sup>1</sup> | 0.056 | 28.833 |
| A0QZJ0 | MSMEG_4042 | Transcriptional regulator, GntR family protein | BACTH <sup>2</sup> | 0.050 | 6.000 |
| A0R616 | MSMEG_6391 | Propionyl-CoA carboxylase beta chain | ClpC1/P2 KD <sup>1</sup> | 0.039 | 9.500 |
| Q3L885 | pks | Polyketide synthase (Type I modular polyketide synthase) | ClpC1/P2 KD <sup>1</sup> | 0.023 | 43.000 |
| A0QQY3 | MSMEG_0918 | Transcriptional regulator, XRE family protein | BACTH <sup>2</sup> | 0.019 | 4.667 |
| A0QSL0 | MSMEG_1516 | Thioredoxin reductase | ClpC1/P2 KD <sup>1</sup> | 0.012 | 9.500 |
| **A0QR26** | MSMEG_0962 | TetR-family protein<br>transcriptional regulator | ClpC1/P2 KD<sup>1</sup><br>BACTH<sup>2</sup> | 0.004 | 42.000 |

<sup>1</sup> (Lunge *et al*, 2020) , <sup>2</sup> (Ziemski et al., 2021)

### Subcomponents of ClpC1 interact with specific cellular proteins

The ClpC1 NTD is a discrete folded module thought to interact with substrates and adaptors, such as the N-end rule adaptor ClpS (22), but is dispensable for hexamer formation, ATP hydrolysis, and unfolding activities. We hypothesized that the NTD and the D1/D2 ATPase core of ClpC1 can bind to different interaction partners. To discriminate between interactions with the NTD and ClpC1 core, we created two truncated ClpC1 constructs: one consisting of only the NTD (ClpC1^NTD^) and another lacking the NTD but possessing the full D1 and D2 core (ClpC1^CORE^) (**Fig 1B**). We repeated our pull-down and proteomics workflow with these constructs.

We observed 243 proteins and 116 proteins that co-immunoprecipitate with ClpC1^NTD^ and ClpC1^CORE^, respectively (**Supplemental Table S1, S2**). Comparing datasets, we found 42 proteins that interact with all core-bearing ClpC1 constructs (ClpC1^WT^, ClpC1^EQ^, and ClpC1^CORE^) (**Fig. 4A**). Notably, 72 out of116 proteins (62%) that interact with the ClpC1^CORE^ were found in at least one other core-bearing datasets, which provides additional confidence that these are *bona fide* interaction partners. 42 proteins were found to interact with all NTD-bearing constructs (ClpC1^WT^, ClpC1^EQ^, and ClpC1^NTD^). Interestingly, only 84 proteins (35%) identified in the ClpC1^NTD^ dataset were observed in at least one of the other NTD-bearing ClpC1^WT^ or ClpC1^EQ^ datasets, suggesting differences in how proteins interact with an isolated monomeric NTD and an NTD in the context of a ClpC1 hexamer.

**Figure 4:**
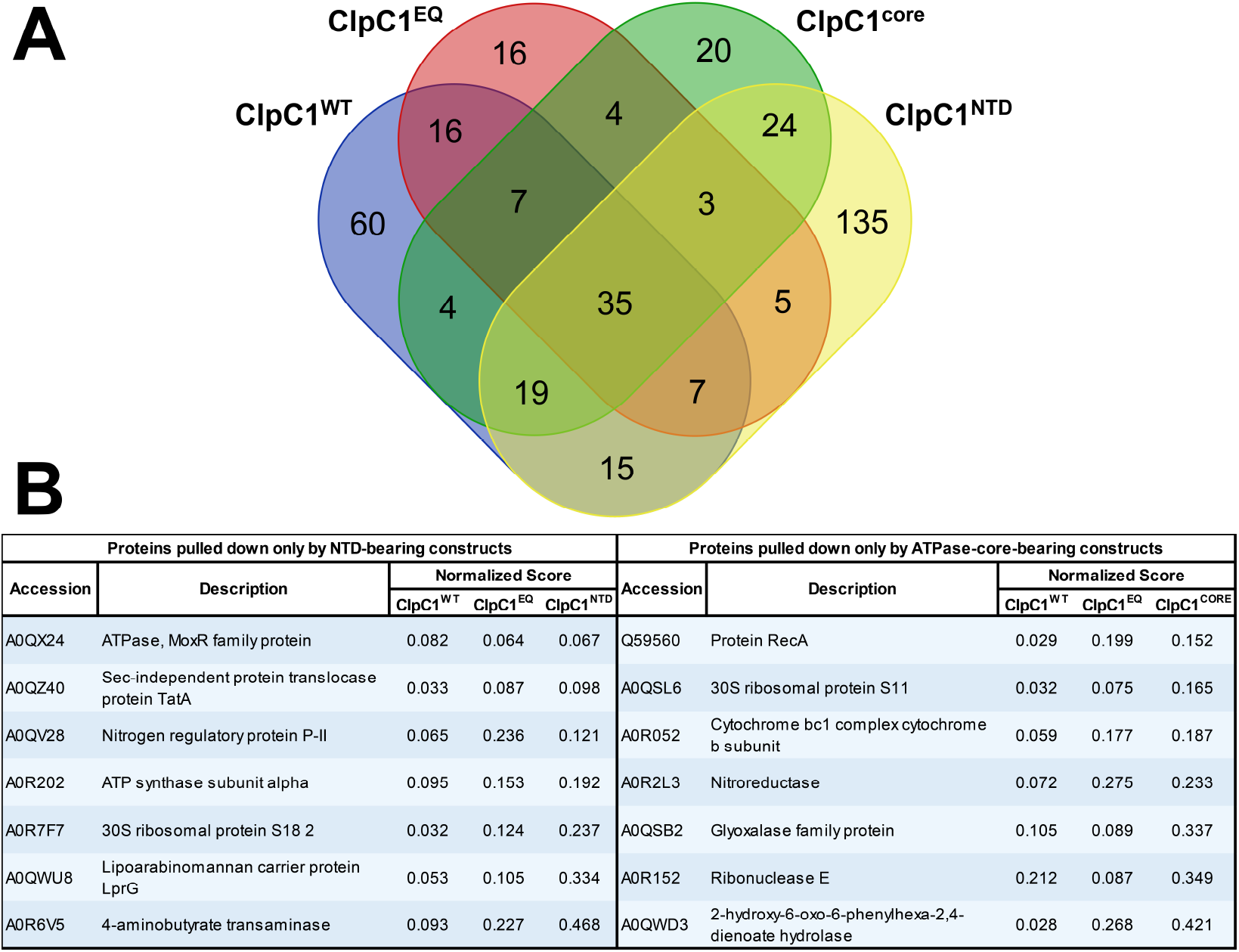
Comparative analysis of ClpC1 interactome based on the different unfoldase components. Venn diagram show overlapping protein targets observed to interact with the ClpC1 constructs bearing the AAA+ ATPase core **(A)** and amino-terminal domain (NTD) **(B)**. Only 7 proteins were observed to interact with NTD alone. Similarly, as shown on the right only 7 proteins were observed interacting with the core-containing constructs, and not in the NTD datasets.

Our analysis identified few proteins that consistently and uniquely interact with NTD-containing or core-containing ClpC1 constructs. Only 7 proteins co-immunoprecipitated with all of NTD-bearing constructs and were absent from the ClpC1^CORE^ dataset. 7 separate proteins interacted with all of core-bearing constructs, but not ClpC1^NTD^ (**Fig. 4B**). Surprisingly, 35 proteins were found in all six datasets, suggesting either that some ClpC1-interacting proteins bind to both NTD and core regions, or that truncated constructs are able to co-assemble with endogenous ClpC1. Overall, our data indicate that many physiological ClpC1 interaction partners exist, and that these bind to multiple regions on the unfoldase.

### Mycobacterial ClpC1 has a diverse and far-reaching interactome

To assess whether specific classes of proteins are preferentially targeted by *Msm* ClpC1, we performed gene ontology (GO) annotation analysis on the interactome using the Blast2GO software suite (60). Gene ontology (GO) annotation analysis was used to assign GO terms for *Mycolicibacterium smegmatis* (strain ATCC 700084 / MC^2^ 155) proteins, based on the closest homologs present in the SwissProt/UniProt database. We found that ClpC1 potentially regulates proteins with a diverse range of cellular functions (**Fig. 5, Table 2**). Of particular interest were protein annotation groups that were highly represented in the full-length ClpC1 interactome. The most represented group of proteins were those associated with nucleic acid-binding and metabolism, including transcriptional regulators (Hup, Rho, RpoC, Rne, RpoA, Pnp, IleS, RpoB, TopA, DnaX, SigA, transcriptional regulators belonging to the MarR, PadR, GntR, XRE, Crp/Fnr, TetR families, among other proteins). Additionally, proteins involved in redox reactions were identified, such as GabD2, IspG, Pks, DapB, among others. Another well-represented group were proteins involved in cellular transport (including SecA1, SecF, Ffh, LprG, TatA). Some of these proteins are transmembrane, but have regions cytoplasmic regions through which proteolytic regulation might occur (71, 72). In addition to nucleic acid metabolism, observed proteins were involved in other forms of metabolism, such as carbohydrate metabolism (GlmS, Eno, SucB, GlpK, SdhA, SdhB, AcnA, Mqo, Kgd), electron transport chain (Ndh, AtpA, AtpC, AtpD, AtpFH), amino acid metabolism (HemL, SerA, GabT, HisG, CarB, GlnA, MSMEG_3973, MSMEG_0688, MSMEI_3879), and lipid metabolism (fatty acid synthase MSMEG_4757, AcpM, acyl-coA synthases MSMEG_0599 and MSMEG_3767). Taken together, these observations suggest that ClpC1 plays a central role in the framework of mycobacterial cellular physiology by potentially regulating metabolic processes and/or ensuring quality control of metabolic enzymes (31).

**Figure 5:**
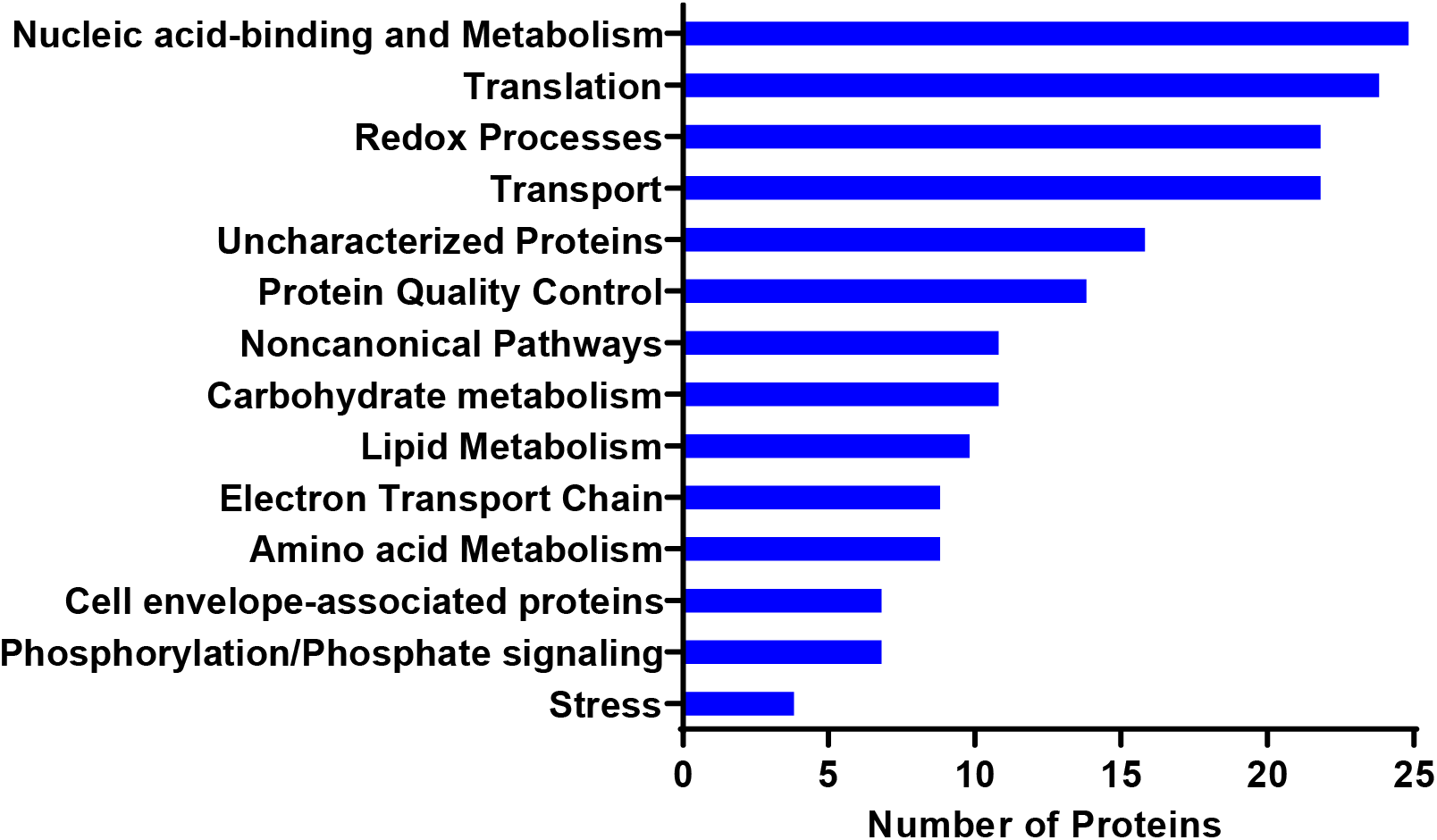
Full-length ClpC1 interacts with diverse groups of proteins in *Mycolicibacterium smegmatis*. Functional categorization of cellular proteins interacting with ClpC1^WT^ and/or ClpC1^EQ^. Shown is the distribution of these proteins by Gene Ontology (GO) annotation based mostly on biological process. Functional classification was performed using Blast2GO, and was based on annotation of closest homologs.

Proteins involved in protein quality control, such as protein folding and unfolding, proteome turnover, and protein homeostasis in general, were also important constituents of the interactome (**Fig. 5, Table 2**). These included chaperone proteins GroL1, GroL2, DnaJ1, DnaJ2, DnaK, Tig, and Clp proteins ClpX and ClpC2 (MSMEG_2792). ClpC1 is thus named because of the existence of the orthologous ClpC2 (MSMEG_2792) which possesses homology to the ClpC1 N-terminal domain, but lacks AAA+ ATPase modules (22, 73). As ClpC1 is itself involved in protein homeostasis and protein fate in general, it is not surprising to see its interaction with these proteins. Also, as noted above, components of the ribosome and other proteins involved in translation were abundant as well, making up ~13% of the total dataset (**Fig. 5**). It is possible that nascent ClpC1 stays in contact with the ribosome during folding, either directly or through indirectly through other elements of folding machinery. Alternatively, ribosomal proteins are commonly observed as contaminants in cell-based mass spectrometry experiments (74).

It is also important to point out that for the proteins specifically interacting with either ClpC1 NTD or core alone (**Fig. 4**), there was some functional diversity. For the NTD-interacting proteins, there were proteins involved in transport (TatA and LprG), phosphorylation/dephosphorylation (MoxR), electron transport chain (AtpA), amino acid metabolism (GabT), noncanonical pathways (MSMEG_2426) and translation (RpsR2). The ClpC1 core-interacting proteins had roles in redox processes (MSMEG_5155 and MSMEG_1417), stress response (RecA), translation (RpsK), electron transport chain (MSMEG_4263), nucleic acid-binding and metabolism (Rne), and noncanonical pathways (MSMEG _2900).

Taken together, we observe a functionally diverse set of cellular proteins that interact with *Msm* ClpC1, revealing a broad diversity of putative substrates and interaction partners of the ClpC1P1P2 protease. This buttresses just how far-reaching its regulatory effects on cellular proteins could be, and may help account for the essentiality of these proteases in mycobacteria (19, 27–31).

### Physiochemical analysis of termini of interacting proteins

Some Clp protease substrates are recognized by short terminal degron sequences of varying length and composition. *Msm* ClpC1 recognizes model substrates bearing a C-terminal ssrA sequence (ADSNQRDYALAA) (13). *Mtb* ClpC1 recognizes Hsp20 via the C-terminal sequence “TQAQRIAITK” (21), and PanD via the C-terminal sequence “NAGELLDPRLGVG” (20). In addition, substrates displaying some hydrophobic N-terminal residues are delivered to ClpC1 by the adaptor ClpS as part of the N-end rule proteolytic pathway (23, 75).

Because the sequence determinants required for recognition by ClpC1 are not well defined, we compared terminal sequences of proteins pulled down by full-length ClpC1 (ClpC1^WT^ or ClpC1^EQ^) to the terminal sequences found across the entire *Msm* proteome (**Fig. 6**). The average charge and hydrophobicity of the first and last 10 amino acids of protein sequences varied widely. We saw no statistically significant differences in these parameters for residues at the N-terminus. However, C-terminal regions of interactome proteins were, on average, less positively charged (closer to neutral) and more hydrophilic than equivalent regions from the full proteome. These differences were statistically significant, although the magnitude of the shift was much smaller than the standard deviation in either dataset. The apparent preference for hydrophilic C-terminal character is surprising, as ATP-dependent proteases are often thought to recognize exposed hydrophobic regions as markers for protein misfolding (8). This analysis suggests either a slight preference for uncharged polar termini, or that polar termini are more exposed to solvent and more available for ClpC1 binding. Additionally, the discrepancy between expected and observed physicochemical parameters might be explained by the presence of ClpC1-interacting proteins that are not proteolytic substrates, and thus may not be subject to the same recognition trends as substrates.

**Figure 6:**
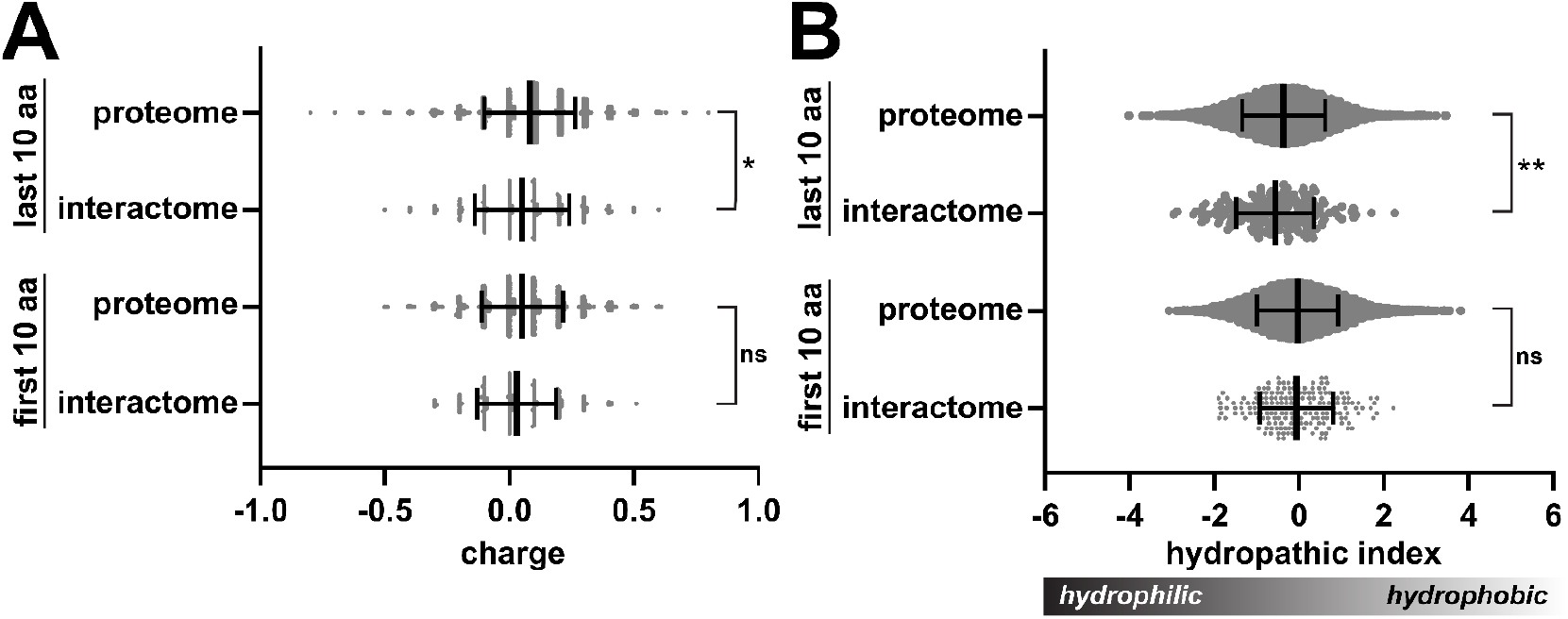
Physicochemical analysis of terminal sequences of full-length ClpC1-interacting proteins. **(A)** Violin plots illustrate average charge of the first (N-terminal) and last (C-terminal) amino acids in the ClpC1 interactome and in the entire *Msm* proteome. Central lines indicate median values, with error bars indicating one standard deviation. **(B)** Mean hydrophobicity of terminal residues in the ClpC1 interactome and control datasets is shown. For both metrics, values plotted are the average residue value per terminus. Significance was assessed by two-tailed Welch’s t-test. * p-value <0.05; ** <0.01; ns not significant.

### ClpC1P1P2 recognizes MSMEI_3879 as a proteolytic substrate via its N-terminal sequence

We selected several hits from our dataset, for which orthologs previously were identified as likely Clp protease interaction partners (**Table 3**) (21, 23), and tested whether these are degraded by ClpC1P1P2 *in vitro*. Candidates included chaperones DnaJ1 and DnaJ2; transcriptional regulators GntR and XRE; magnesium chelatase; and the apparent pseudogene product MSMEI_3879. Of the proteins tested, only MSMEI_3879 was degraded by ClpC1P1P2, with ~75% of the substrate hydrolyzed within 40 min (**Fig. 7A, B**). These *in vitro* results provide preliminary evidence of the substrate status of MSMEI_3879.

**Figure 7.**
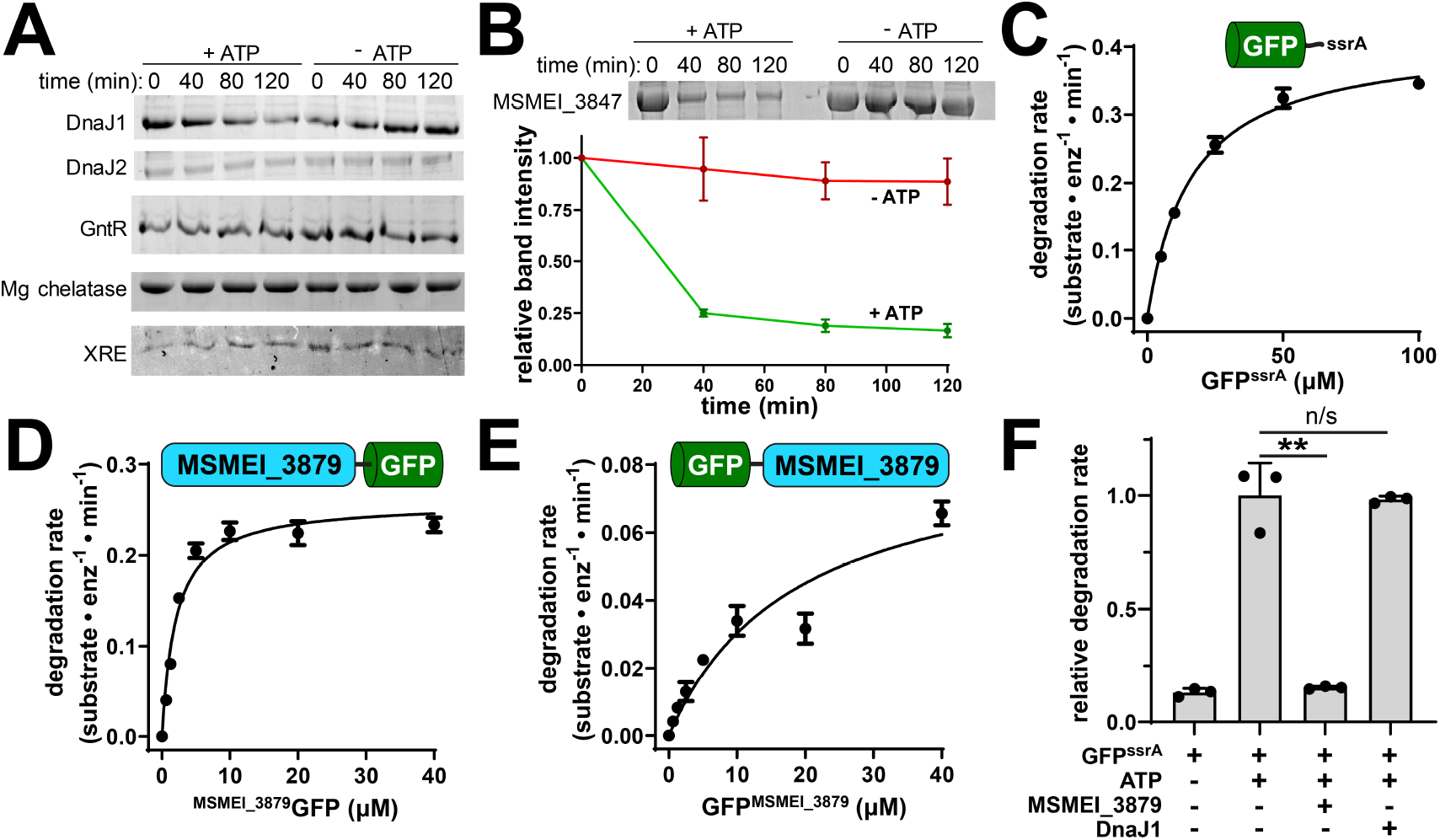
MSMEI_3879 is a substrate of *Msm* ClpC1. **(A)** *In vitro* degradation assays of chaperones DnaJ1, DnaJ2; transcriptional regulators GntR, magnesium chelatase, and XRE by 1 µM ClpC1 and 1 µM ClpP1P2, monitored by SDS/PAGE. **(B)** ATP-dependent degradation of 10 µM MSMEI_3879 by ClpC1P1P2 was observed *in vitro* by SDS-PAGE, with gel densitometry of three replicate assays shown. **(C)** Michaelis-Menten analysis of GFP^ssrA^ proteolysis by 1 µM ClpC1P1P2, as function of substrate concentration, reveals a *k_cat_* of 0.41 ± 0.02 substrate•min^−1^•enz^−1^ and a *K_M_* of 16.04 ± 1.28 µM. **(C)** ^MSMEI_3879^GFP was degraded by 1 µM ClpC1P1P2 with a *k_cat_* of 0.26 ± 0.01 substrate•min^−1^•enz^−1^ and *K_M_* of 2.06 ± 0.43 µM. **(E)** GFP^MSMEI_3879^ was degraded with a *k_cat_* of 0.09 ± 0.02 substrate•min^−1^•enz^−1^ and a *K_M_* of 18.8 ± 9.6 µM. **(F)** Degradation 10 µM GFP^ssrA^ by 1 µM ClpC1P1P2 was monitored in the presence and absence of ATP, 10 µM MSMEI_3879, or 10 µM DnaJ1. MSMEI_3879 reduces GFP^ssrA^ degradation to the level observed in the absence of ATP. DnaJ1 does not significantly alter the rate of GFP^ssrA^ proteolysis. Values are averages of three replicates (N = 3) ± 1 SD. p-values were calculated by unpaired two-tailed Student’s t-test. **: *p* value < 0.01; n.s.: not significant.

**Supplemental Figure S1:**
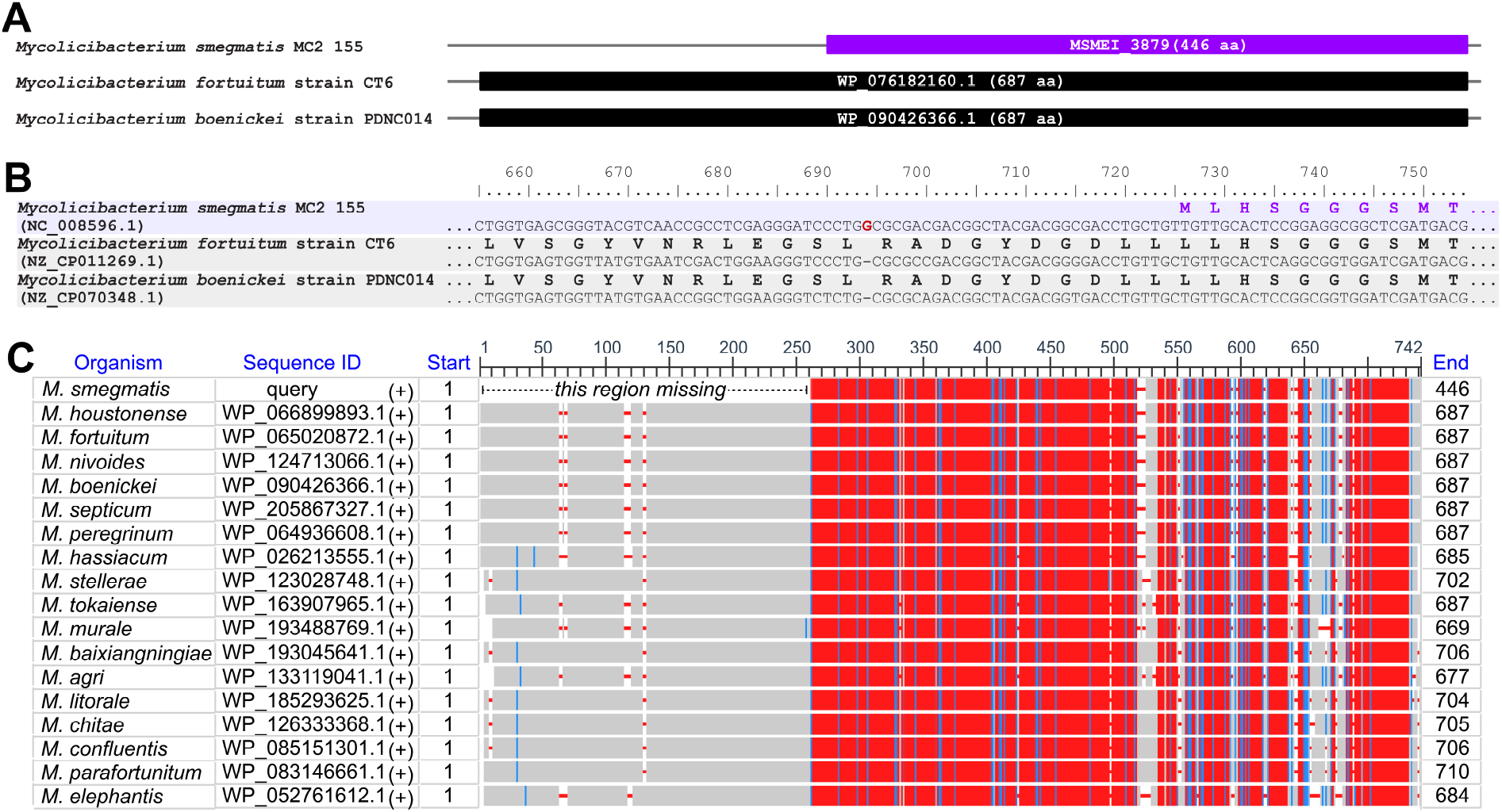
Comparison of MSMEI_3879 to full-length homologs. **(A)** Genetic context of *M. smegmatis* MSMEI_3879 compared to the equivalent genomic region in the indicated *Mycolicibacterium* species. **(B)** Sections of genomic sequence show the beginning of the disrupted *M. smegmatis* hydantoinase/oxoprolinase gene and the analogous regions in the intact genes of *M. fortuitum* and *M. boenickei*. Numbering is given from the beginning of the normal intact open reading frame. Amino acid translations of MSMEI_3879 (purple) and hydantoinase/oxoprolinase orthologs (black) are shown. A single nucleotide insertion in the *M. smegmatis* genome (red) disrupts the gene. **(C)** NCBI protein BLAST was used to align MSMEI_3879 (top) with full-length hydantoinase/oxoprolinase orthologs from select *Mycolicibacterium* species. The upstream region missing in MSMEI_3879 is indicated.

The MSMEI_3879 locus encodes a 446 amino acid product with homology to ATP-hydrolyzing hydantoinase/oxoprolinase enzymes, which catalyze ring-opening reactions on lactam substrates (76–78). Orthologs with >90% sequence identity to MSMEI_3879 are present in members of the genus *Mycolicibacterium*, but absent in other groups of the family *Mycobacteriaceae*, including *M. tuberculosis* (79). Notably, *Mycolicibacterium* orthologs of MSMEI_3879 are generally longer, at about 690 amino acids. Comparison of the *M. smegmatis* genome to those of other *Mycolicibacterium* species reveals a frameshift caused by a single nucleotide insertion in codon 233 (**Fig. S1**). The MSMEI_3879 open reading frame arises from an alternative start site at position 242 (with respect to the typical *Mycolicibacterium* start codon), and the resulting polypeptide corresponds to only the latter two-thirds of orthologous hydantoinase/oxoprolinase enzymes. (There is no MSMEG annotation that directly corresponds to MSMEI_3879. The closest analog, MSMEG_3974, describes the entire frameshifted locus.) While MSMEI_3879 is annotated as 5-oxoprolinase, truncation likely abolishes its enzymatic activity. In spite of this, MSMEI_3879 expressed well in *E. coli* and behaved well *in vitro*.

Many Clp protease substrates are recognized by short terminal degron sequences (13, 20, 21, 23). To test whether ClpC1 recognizes MSMEI_3879 by a particular terminus, we engineered GFP constructs with MSMEI_3879 fused to either the N- or C-terminus, and assayed proteolysis by ClpC1P1P2. Michaelis-Menten analysis of the resulting degradation rates showed that the construct with MSMEI_3879 at the N-terminus (^MSMEI_3879^GFP) is degraded with *k_cat_* ≈ 0.26 sub•min^−1^•enz^−1^ and *K_M_* ≈ 2 µM (**Fig. 7C**). For comparison, under the same assay conditions, GFP with a *Msm* ssrA tag (GFP^ssrA^) was degraded faster (*k_cat_* ≈ 0.4 sub•min^−1^•enz^−1^) but with a much higher *K_M_* ≈ 16 µM (**Fig. 7D**), similar to the *K_M_* reported for degradation of GFP^ssrA^ by *Mtb* ClpC1 (23). A construct carrying MSMEI_3879 at the C-terminus (GFP^MSMEI_3879^) was degraded at a substantially slower rate (*k_cat_* ≈ 0.09 sub•min^−1^•enz^−1^) and higher *K_M_* (~19 µM), compared to ^MSMEI_3879^GFP (**Fig. 7E**). The 9-fold lower *K_M_* of ^MSMEI_3879^GFP suggests that the N-terminus of MSMEI_3879 contributes to efficient degradation by ClpC1, but is not the sole determinant of recognition. It is likely that the genetic truncation of MSMEI_3879 creates disordered or unstructured matter at the N-terminus that contributes to robust ClpC1 recognition.

Given the lower *K_M_* for degradation of MSMEI_3879 compared to GFP^ssrA^, we tested whether untagged MSMEI_3879 can compete with GFP^ssrA^ degradation and found that equimolar MSMEI_3879 effectively blocks GFP^ssrA^ proteolysis (**Fig. 7F**). Inhibition was a specific feature of MSMEI_3879, as DnaJ1, a non-substrate protein from our interactome dataset (**Fig. 7A**), had no effect on degradation (**Fig. 7F**). Taken together, these data indicate MSMEI_3879 is a novel and robustly-degraded ClpC1P1P2 substrate, and that its N-terminal sequence contributes to its recognition.

### Chemoinhibition of ClpC1 blocks MSMEI_3879 degradation

We sought to assess whether MSMEI_3879 constructs would have utility as reporters for small molecule disruption of ClpC1. We used two well-characterized mycobacterial ClpC1 dysregulators, ecumicin (ECU) and rufomycin (RUF), which bind the ClpC1 NTD and dysregulate the ability of ClpC1 to degrade protein substrates (80–82). We assayed proteolysis of ^MSMEI_3879^GFP and GFP^ssrA^ by ClpC1P1P2 *in vitro* in the presence and absence of 10 µM of each compound. Compared to the untreated control, both ECU and RUF reduced the rate of ^MSMEI_3879^GFP degradation by about 60% (**Fig. 8A**). Inhibition of GFP^ssrA^ degradation was also observed, but with different magnitude for each compound: ECU inhibited by ~80% while RUF inhibited by ~45% (**Fig. 8B**). Taken together, these results demonstrate that known ClpC1 dysregulators inhibit proteolysis of MSMEI_3879 by ClpC1P1P2.

**Figure 8:**
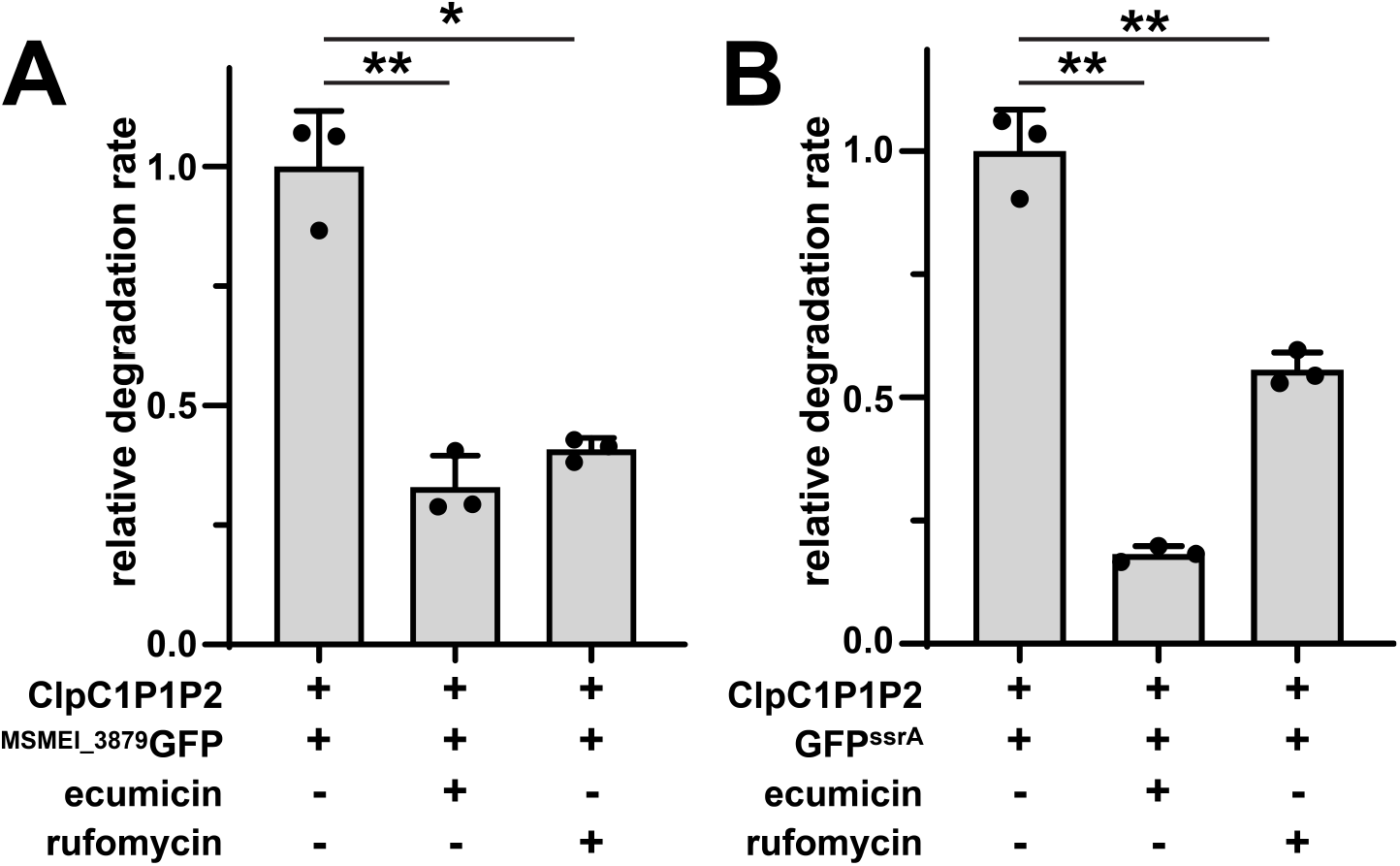
Effect of ClpC1-targeting antibiotics on proteolysis by ClpC1P1P2. Inclusion of 10 µM ECU or RUF inhibits degradation of 10 µM **(A)** ^MSMEI_3879^GFP and **(B)** GFP^ssrA^ by 1 µM ClpC1P1P2, relative to a non-treatment control. Values are averages of three replicates (N = 3) ± 1 SD. p-values were calculated by unpaired two-tailed Student’s t-test. * and ** represent *p* values < 0.05 and 0.01, respectively.

## Discussion

It is now well established that Clp protease components, including ClpC1, are essential for mycobacterial viability (19, 27–31), and these enzymes have consequently emerged as promising antibiotic targets. Here, we sought to better understand the physiological roles of ClpC1P1P2 by identifying partners that interact with full-length and truncated ClpC1 constructs. In total, we found 370 unique cellular proteins that interact with one or more ClpC1 construct. The diversity of the identified ClpC1 interactome is consistent with a multifaceted role for this enzyme in mycobacterial physiology. Moreover, our data aligns with recent explorations of the *Mtb* ClpC1 interactome/degradome (21, 23), as 30 proteins found here possess orthologs identified in those screens.

While ClpC1 likely participates in multiple proteolytic programs in mycobacteria, we note that not all interacting proteins identified here are substrates. Of the six proteins we tested directly, only one, MSMEI_3879, proved to be a *bona fide* substrate *in vitro* (**Fig. 7**). Non-substrate interaction partners may play a role in regulating ClpC1 activity, may selectively interact with protease-deficient higher-order ClpC1 oligomers (25, 83, 84), or may simply bind non-specifically under our capture conditions. It is also notable that capture experiments utilizing ClpC1^EQ^, a construct carrying Walker B mutations expected to stabilize interactions with proteolytic substrates (56, 68–70), yielded fewer identified interaction partners (93 proteins) than experiments with ClpC1^WT^ (163 proteins) (**Fig. 3; Tables 1 and 2; Supplemental Table S2**). This suggests that many of these interaction partners are not engaged as substrates, and instead adopt different modes of interaction that are dependent on nucleotide binding, ATP hydrolysis, and perhaps oligomeric state. A diverse set of dynamic interaction partners would plausibly allow mycobacterial cells to nimbly modulate ClpC1 activity across different growth conditions.

ClpC1 is composed of a globular N-terminal domain that binds to substrates and adaptors, and an ATPase core consisting of two AAA+ modules that carry out chemomechanical substrate unfolding and translocation (8, 10, 85–87). Surprisingly, our studies identified few interaction partners that bind exclusively to either the NTD or the ATPase core. This suggests that many ClpC1-interacting proteins make multivalent binding to both components. Alternatively, this could result from truncated constructs (ClpC1^NTD^ and ClpC1^CORE^) assembling with endogenous ClpC1 in the cell. Regardless of the mechanism involved, this illustrates that ClpC1 acts through the cooperation of its constituent parts.

Importantly, our interactome analysis led us to identify MSMEI_3879 as a *bona fide* ClpC1P1P2 substrate (**Fig. 7**). This protein joins a short list of verified mycobacterial Clp protease substrates (20, 21, 23, 24). MSMEI_3879 is degraded relatively efficiently: a GFP construct carrying an N-terminal MSMEI_3879 fusion was degraded with a *K_M_* ~7-fold lower than the model substrate GFP^ssrA^ (13, 17). As expected for this lower *K_M_*, degradation of GFP^ssrA^ was effectively blocked by an equimolar amount of MSMEI_3879 (**Fig. 7F**). Our experiments indicate that recognition by ClpC1 requires an exposed MSMEI_3879 N-terminus. This likely correlates with the fact that MSMEI_3879 is a truncated gene product, arising from a frameshift mutation in the typical mycobacterial hydantoinase/oxoprolinase locus that eliminates the first ~240 residues (**Supplemental Figure S1**). Recognition by ClpC1 may occur as a consequence of this truncation, via unstructured or poorly folded regions present at the MSMEI_3879 N-terminus. Indeed, ClpC1 has been proposed to selectively recognize some substrates via disordered termini (21). It is unclear whether MSMEI_3879 carries out any significant enzymatic function, or whether its proteolysis by ClpC1 plays any role in *M. smegmatis* physiology. Regardless, MSMEI_3879-based substrates may serve as useful tools for probing ClpC1P1P2 activity and dysregulation. As a proof-of-concept, we show that known ClpC1-targeting antimicrobials inhibit ^MSMEI_3879^GFP proteolysis *in vitro* (**Fig. 8**). Similar substrates could serve as the basis for cell-based screens to identify novel ClpC1-targeting antibiotics.

## Acknowledgements

We gratefully thank R. Neunuebel and V. Parashar for the use of instrumentation. We thank Papa-Nii Asare-Okai for technical support.

